## Supplemental figures for "Bone marrow adipocytes drive the development of tissue invasive Ly6C^high^ monocytes during obesity"

**Figure S6: Flow cytometry gating strategy for identification of progenitor cells within Ly6C$^{low}$ and Ly6C$^{high}$ monocyte subsets.** To prevent contamination of blood in BM cells, mice (n=3 individual mice) were placed under deep anesthesia using isoflurane and underwent intra-cardiac perfusion with 10 mL of cold PBS, before sacrifice. Femurs were surgically removed and crushed with a mortar and pestle and bone marrow cells suspended in PBS were filtered through a 40-µm-nylon mesh. Cells were pelleted at 300 × g, 5 min and resuspended in 500 µl FACS Buffer. 100 µl was aliquoted and stained with the antibody cocktail for 30 min. Antibodies for CD117, CD16/32 and CD34, were used at 1:50 dilution, all others at 1:100 dilution. Samples were fixed (BD, Cytofix) before being run on a BD LSR Fortessa cytometer. [A] Gating strategy for Ly6C$^{low}$ and Ly6C$^{high}$. [B] Gating strategy for progenitor cell identification in the Ly6C$^{high}$ monocytes. [C] Gating strategy for progenitor cell identification in the Ly6C$^{low}$ monocytes.

### Figure S1

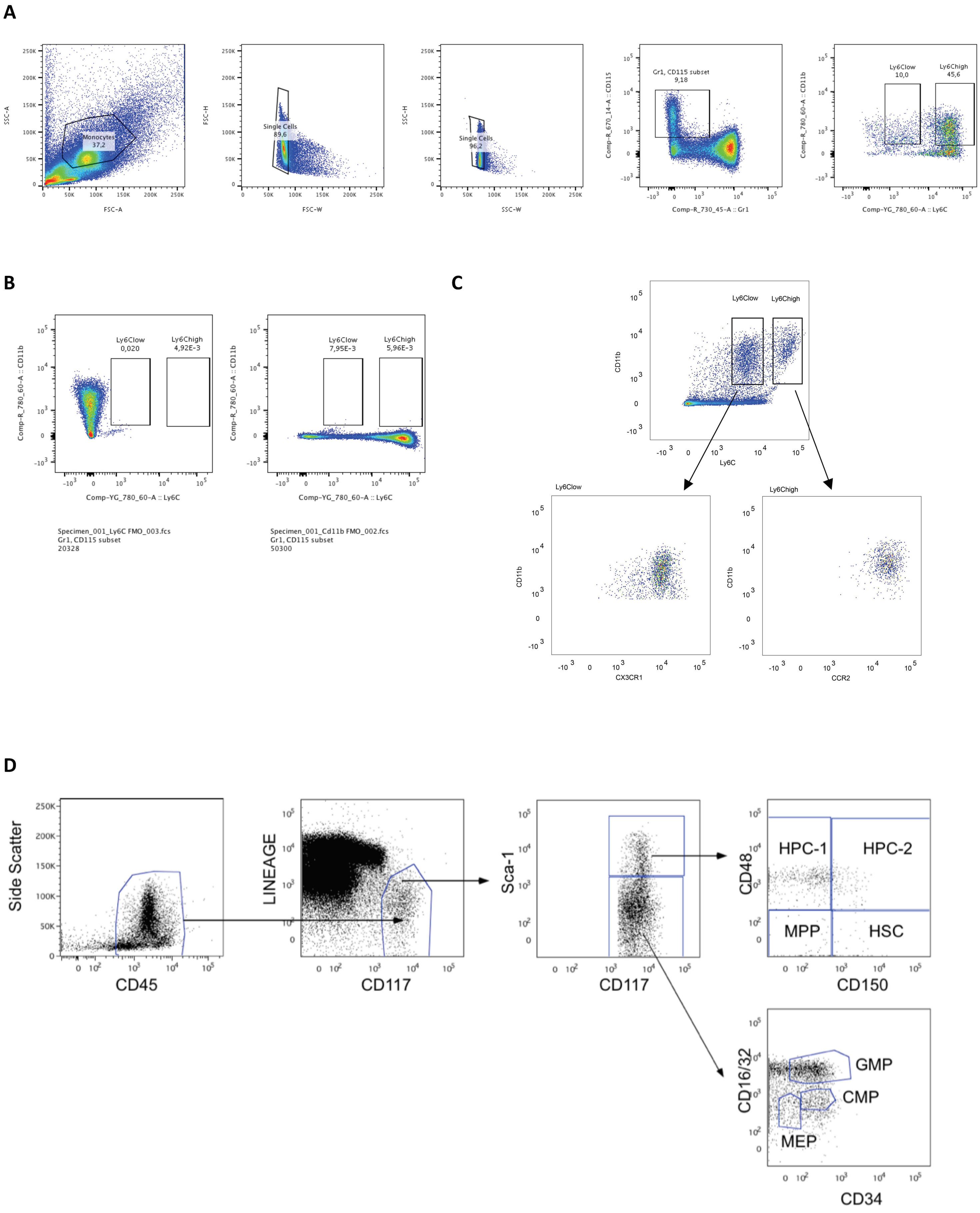
A
Monocytes 37,2
SSC-A
FSC-A
Single Cells 89,6
FSC-H
FSC-W
Single Cells 96,2
SSC-W
Gr1, CD115 subset 9,18
Comp-R_670_14-A :: CD115
Comp-R_730_45-A :: Gr1
Ly6Clow 10,0
Ly6Chigh 45,6
Comp-R_780_60-A :: CD11b
Comp-YG_780_60-A :: Ly6C
B
Ly6Clow 0,020
Ly6Chigh 4,92E-3
Specimen_001_Ly6C FMO_003.fcs
Gr1, CD115 subset
20328
Ly6Clow 7,95E-3
Ly6Chigh 5,96E-3
Specimen_001_Cd11b FMO_002.fcs
Gr1, CD115 subset
50300
C
Ly6Clow
Ly6Chigh
CD11b
Ly6C
CX3CR1
CCR2
D
Side Scatter
CD45
LINEAGE
CD117
Sca-1
CD48
CD150
HPC-1
HPC-2
MPP
HSC
CD16/32
CD34
GMP
CMP
MEP

**Figure S2**

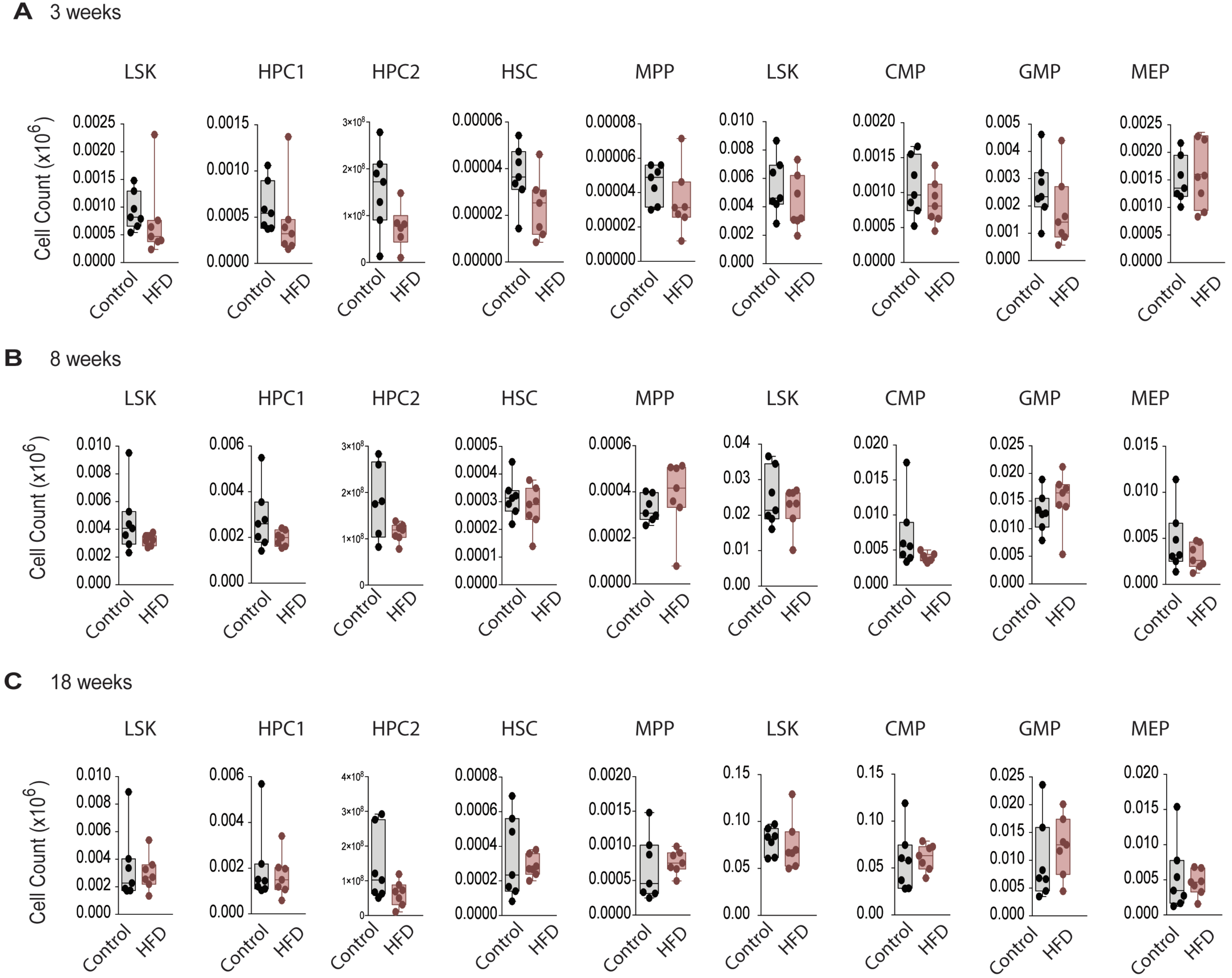

A 3 weeks
B 8 weeks
C 18 weeks
LSK
HPC1
HPC2
HSC
MPP
CMP
GMP
MEP
Cell Count (x10^6)
Control
HFD

**Figure S3**

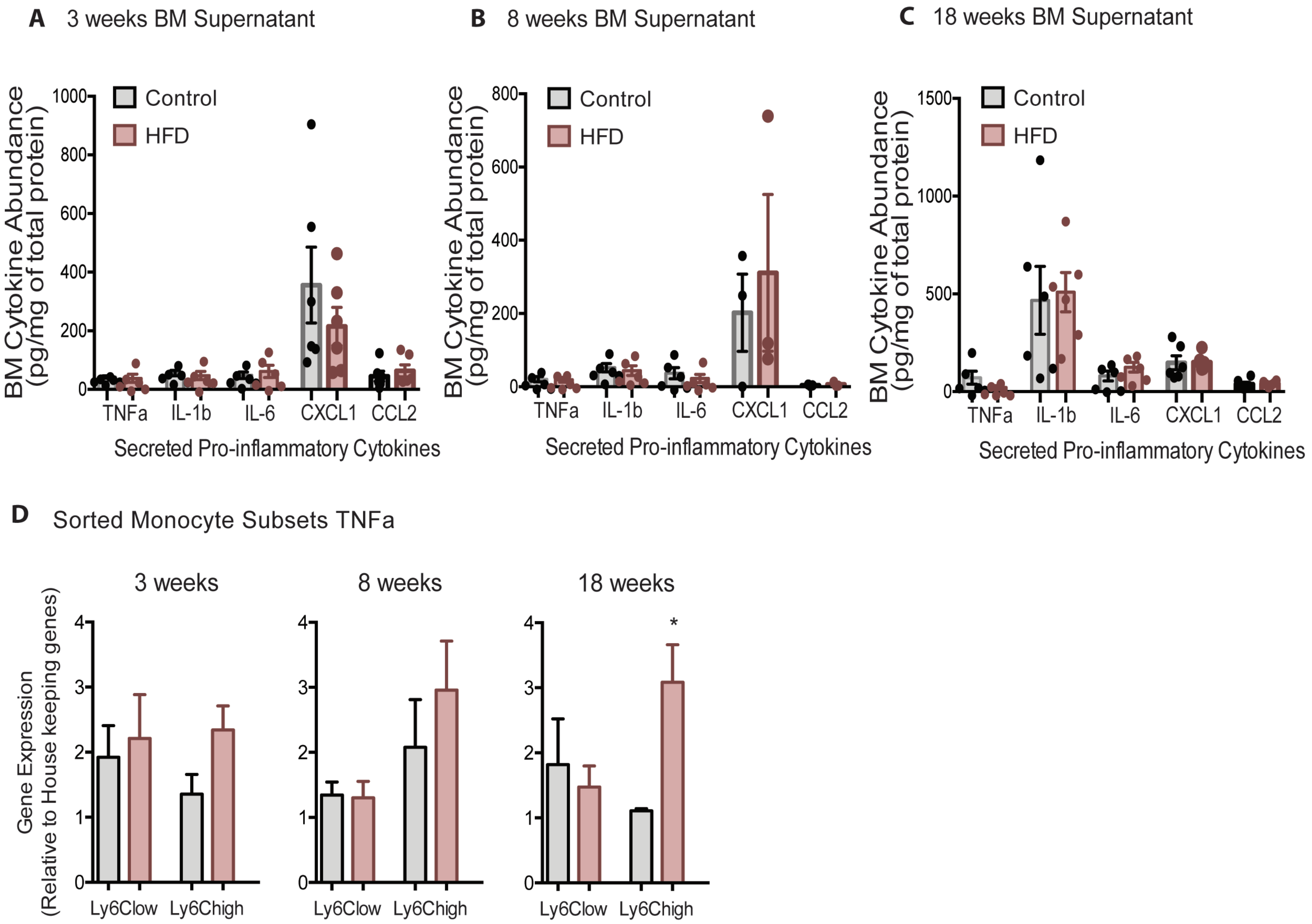
A
3 weeks BM Supernatant
B
8 weeks BM Supernatant
C
18 weeks BM Supernatant
BM Cytokine Abundance
(pg/mg of total protein)
Control
HFD
1000
800
600
400
200
0
1500
500
TNFa
IL-1b
IL-6
CXCL1
CCL2
Secreted Pro-inflammatory Cytokines
D
Sorted Monocyte Subsets TNFa
3 weeks
8 weeks
18 weeks
Gene Expression
(Relative to House keeping genes)
4
3
2
1
0
*
Ly6Clow
Ly6Chigh

### Figure S4

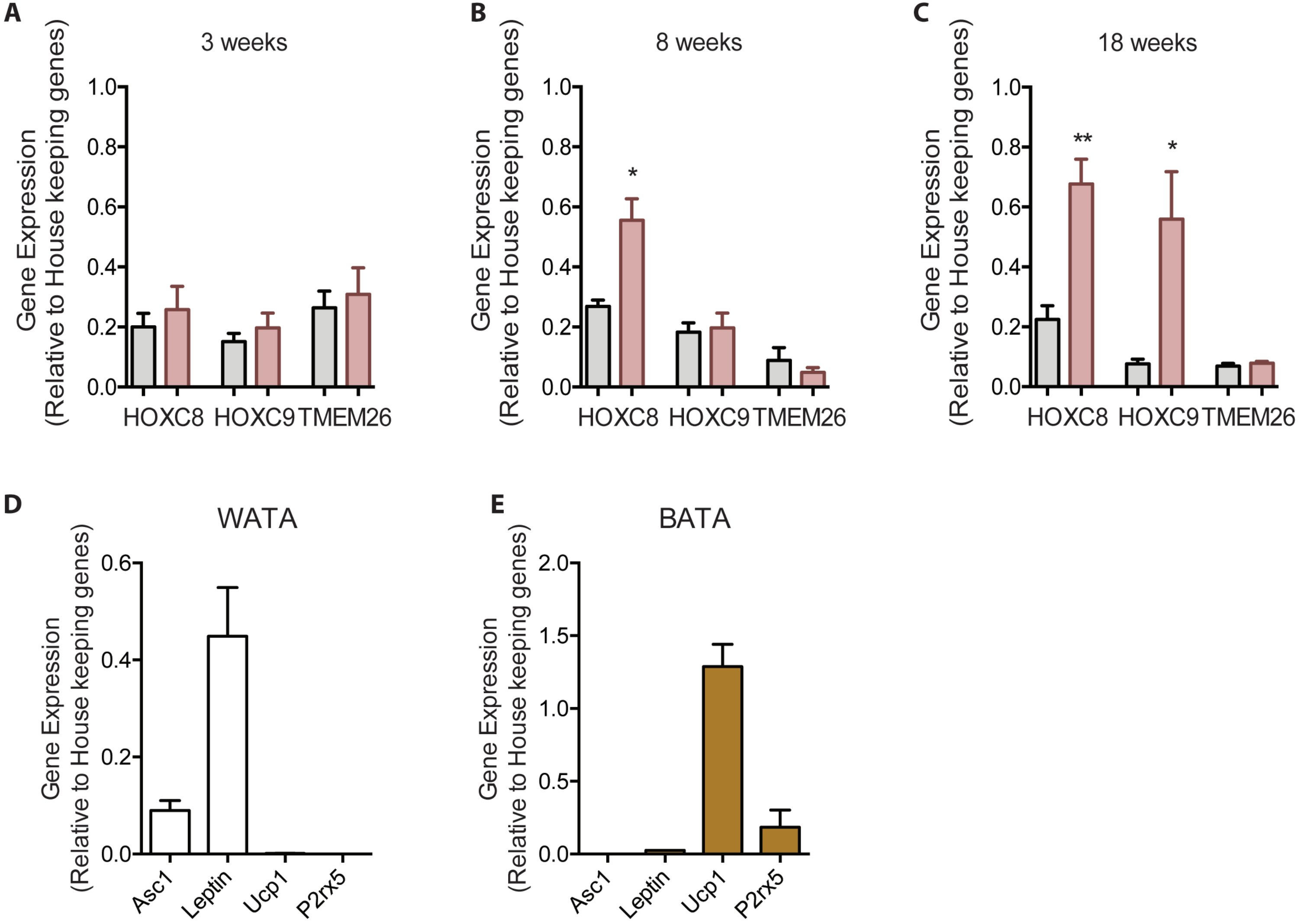

A
3 weeks
Gene Expression (Relative to House keeping genes)
1.0
0.8
0.6
0.4
0.2
0.0
HOXC8
HOXC9
TMEM26
B
8 weeks
*
C
18 weeks
**
*
D
WATA
0.6
0.4
0.2
0.0
Asc1
Leptin
Ucp1
P2rx5
E
BATA
2.0
1.5
1.0
0.5
0.0

**Figure S5**

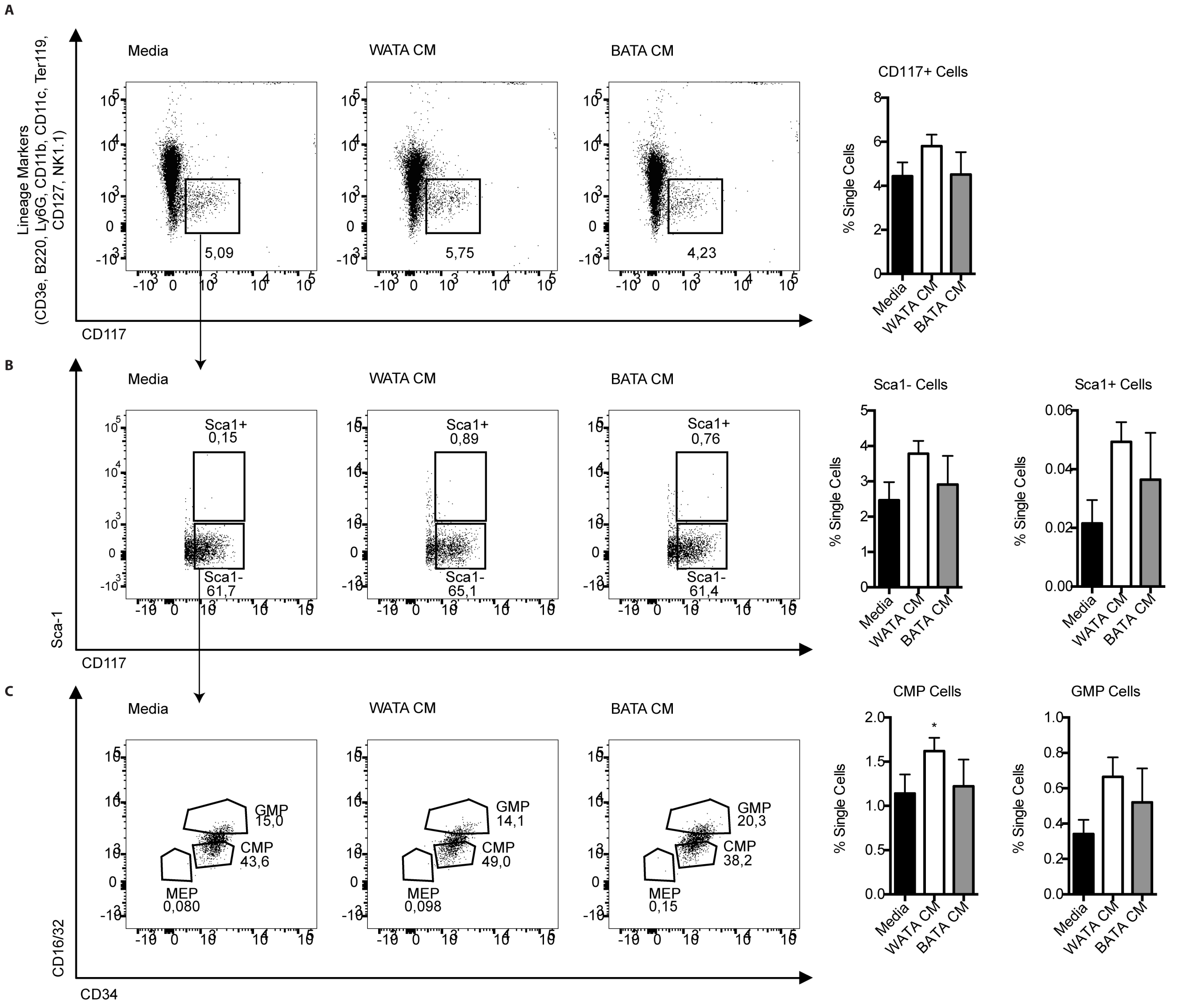
A
Media
WATA CM
BATA CM
5,09
5,75
4,23
Lineage Markers (CD3e, B220, Ly6G, CD11b, CD11c, Ter119, CD127, NK1.1)
CD117
CD117+ Cells
% Single Cells
Media
WATA CM
BATA CM
B
Media
WATA CM
BATA CM
Sca1+
0,15
0,89
0,76
Sca1-
61,7
65,1
61,4
Sca-1
CD117
Sca1- Cells
Sca1+ Cells
% Single Cells
C
Media
WATA CM
BATA CM
GMP
15,0
14,1
20,3
CMP
43,6
49,0
38,2
MEP
0,080
0,098
0,15
CD16/32
CD34
CMP Cells
GMP Cells
% Single Cells

### Figure S6

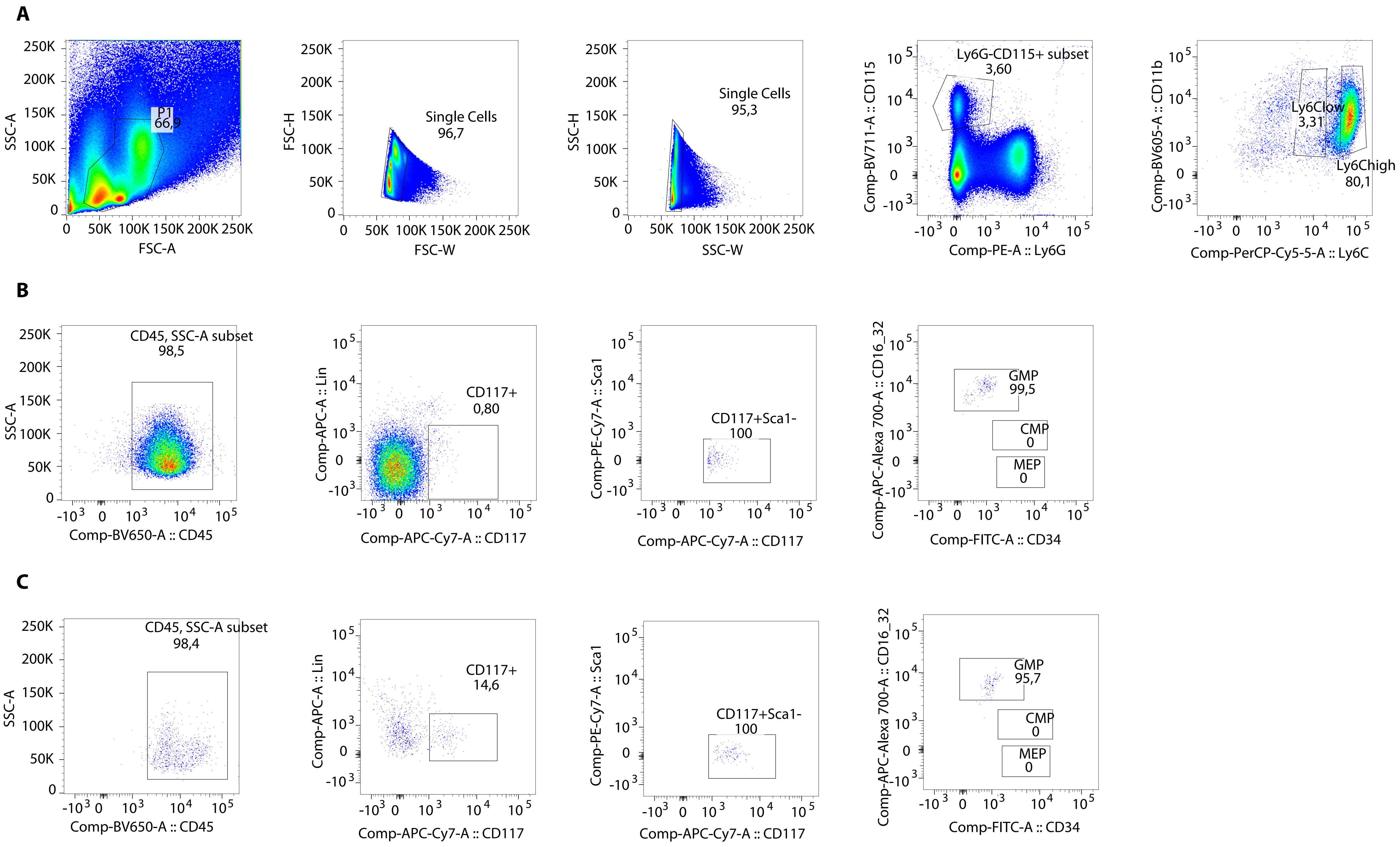

A
P1
66,9
SSC-A
FSC-A
Single Cells
96,7
FSC-H
FSC-W
Single Cells
95,3
SSC-H
SSC-W
Ly6G-CD115+ subset
3,60
Comp-BV711-A :: CD115
Comp-PE-A :: Ly6G
Ly6Clow
3,31
Ly6Chigh
80,1
Comp-BV605-A :: CD11b
Comp-PerCP-Cy5-5-A :: Ly6C
B
CD45, SSC-A subset
98,5
SSC-A
Comp-BV650-A :: CD45
CD117+
0,80
Comp-APC-A :: Lin
Comp-APC-Cy7-A :: CD117
CD117+Sca1-
100
Comp-PE-Cy7-A :: Sca1
Comp-APC-Cy7-A :: CD117
GMP
99,5
CMP
0
MEP
0
Comp-APC-Alexa 700-A :: CD16_32
Comp-FITC-A :: CD34
C
CD45, SSC-A subset
98,4
SSC-A
Comp-BV650-A :: CD45
CD117+
14,6
Comp-APC-A :: Lin
Comp-APC-Cy7-A :: CD117
CD117+Sca1-
100
Comp-PE-Cy7-A :: Sca1
Comp-APC-Cy7-A :: CD117
GMP
95,7
CMP
0
MEP
0
Comp-APC-Alexa 700-A :: CD16_32
Comp-FITC-A :: CD34
